# Humanized NBSGW PDX Models of Disseminated Ovarian Cancer Recapitulate Key Aspects of the Tumor Immune Environment within the Peritoneal Cavity

**DOI:** 10.1101/2022.07.01.498335

**Authors:** Mara P. Steinkamp, Irina Lagutina, Kathryn J. Brayer, Fred Schultz, Danielle Burke, Vernon S. Pankratz, Sarah F. Adams, Laurie G. Hudson, Scott A. Ness, Angela Wandinger-Ness

## Abstract

The importance of the immune microenvironment in ovarian cancer progression, metastasis, and response to therapies has become increasingly clear, especially with the new emphasis on immunotherapies. In order to leverage the power of patient-derived xenograft (PDX) models within a humanized immune microenvironment, three ovarian cancer PDX were grown in humanized NBSGW mice engrafted with human CD34+ cord blood-derived hematopoietic stem cells. Analysis of cytokine levels in the ascites fluid and infiltrating immune cells in the tumors demonstrated that these humanized PDX (huPDX) established an immune tumor microenvironment similar to what has been reported for ovarian cancer patients. The lack of human myeloid cell differentiation has been a major setback for humanized mouse models, but our analysis shows that PDX engraftment increases the human myeloid population in the peripheral blood. Analysis of cytokines within the ascites fluid of huPDX revealed high levels of human M-CSF, a key myeloid differentiation factor as well as other elevated cytokines that have previously been identified in ovarian cancer patient ascites fluid including those involved in immune cell differentiation and recruitment. Human tumor-associated macrophages and tumor-infiltrating lymphocytes were detected within the tumors of humanized mice, demonstrating immune cell recruitment to tumors. Comparison of the three huPDX revealed certain differences in cytokine signatures and in the extent of immune cell recruitment. Our studies show that huNBSGW PDX models reconstitute important aspects of the ovarian cancer immune tumor microenvironment making this a superior approach for therapeutic trials.

## INTRODUCTION

The search for novel treatment regimens for ovarian cancer would benefit from preclinical models that represent the genetic heterogeneity of the patient population, while recapitulating the unique ovarian cancer tumor microenvironment including the tumor-promoting immune cell populations. Previous studies have shown that patient derived xenograft (PDX) models retain the genetic heterogeneity of the primary tumor and recapitulate the original tumor morphology^1-4^. Use of multiple PDX models is advantageous for sampling the heterogeneity across the patient population and increases the translational potential of a given intervention^4-6^. However, PDX models growing in immunocompromised mice lack the immune components of the tumor microenvironment.

Irradiated, immunocompromised NSG mice engrafted with CD34+ hematopoietic stem cells (HSCs) yield primarily human B and T cells and are the standard humanized mouse model for oncology studies. However, this model has a low representation of myeloid populations, particularly macrophages^7^. The lack of myeloid cells is a critical shortcoming in the standard humanized NSG model particularly for ovarian cancer research. Ovarian cancer cells produce cytokines that recruit macrophages to tumors and skew macrophage polarization towards an anti-inflammatory state^8^. In turn, tumor associated macrophages (TAMs) release a range of factors such as epidermal growth factor (EGF) that stimulate cancer cell proliferation and metastasis^9^. TAMs make up 30%-50% of the cells found in malignant ascites^10^. Therefore, it is critical that TAMs be present in humanized ovarian cancer PDX models.

With the recognition that myeloid cells are critical in tumor biology and immunity, attempts have been made to enrich myeloid cells in humanized models. New strategies make use of transgenic mice that constitutively express human cytokines to improve myeloid differentiation. The NSG-SGM3 transgenic model that constitutively expresses human GM-CSF, has a larger number of myeloid progenitor cells in the bone marrow, but does not show an increase in differentiated myeloid populations in the peripheral blood^11^. The MISTRG mouse model that constitutively expresses human M-CSF has higher numbers of myeloid cells in the blood, but rapidly develops anemia, which can reduce the window for performing cancer studies^12, 13^. Thus, these strategies are not yet ideal.

Recent studies examining the anti-cancer effects of human immune cells in ovarian cancer xenograft models have adoptively transferred subsets of human immune cells including allogeneic NK-cells^14^, allogeneic tumor-primed human T cells^15^, and autologous mature T cells from patient tumors^16^. None of these models reconstitutes the full complement of human immune cells within the tumor microenvironment. One study engrafted autologous PBMCs or tumor-associated leukocytes into a subcutaneous ovarian cancer PDX model^17^. However, this model requires the isolation of large numbers of autologous tumor-associated leukocytes and does not translate easily to large-scale therapeutic studies. Furthermore, subcutaneous tumors do not reflect the peritoneal ovarian tumor environment. Our aim was to develop a disseminated ovarian cancer PDX model with a more complete humanized immune system to better represent the unique immune microenvironment within the peritoneal cavity.

Here, we present a strategy for ovarian cancer growth in humanized mice that results in myeloid engraftment, differentiation, and tumor infiltration. We establish disseminated ovarian cancer PDX models in humanized NBSGW mice engrafted with human cord-blood derived CD34+ HSCs. The NBSGW strain on the NSG background carries a *c-kit* mutation that weakens the ability of mouse HSCs to compete with engrafted human cells. Humanized NBSGW demonstrate efficient humanization and increased CD33+ myeloid progenitor cells in the bone marrow and CD11b+ myeloid cells and CD1a+ dendritic cells within the spleen without the need for pre-engraftment irradiation^**18**, **19**^. By engrafting ovarian cancer PDX models into humanized NBSGW mice, we can take advantage of the genetic heterogeneity of the PDX models, while restoring the complex interactions between human cancer cells and human immune cells. Cytokine profiling and evaluation of tumor infiltrating immune cells in multiple huPDX models demonstrated a reconstitution of the peritoneal tumor immune environment in these models.

## MATERIALS AND METHODS

### Banking patient ovarian cancer spheroids and establishing PDX models from malignant ascites

Malignant ascites from consenting ovarian cancer patients were collected during cytoreductive surgery at the University of New Mexico Comprehensive Cancer Center (UNMCCC). Acquisition of patient samples was approved by the UNM Health Science Center Institutional Review Board (protocol #INST1509). Ascites samples were centrifuged and cell-free ascites fluid was stored at -80°C for cytokine analysis. After RBC lysis using Ammonium Chloride Solution, cancer spheroids were isolated and 20 × 10^6^ ovarian cancer cells were injected into the peritoneal cavity of NSG mice to establish orthotopic PDX models of disseminated ovarian cancer. Mice were euthanized at a humane endpoint when mice exhibited abdominal distention from ascites accumulation or signs of wasting. Solid tumors and ascites fluid were collected. PDX samples were cryopreserved in 95% FBS/ 5% DMSO or injected into new NSG mice for passaging. All mouse procedures were approved by the UNM Animal Care and Use Committee (Protocol#18-200722-HSC), in accordance with NIH guidelines for the Care and Use of Experimental Animals.

### RNA isolation and RNA-seq of ovarian cancer samples

RNA-seq analysis was performed on primary ovarian cancer solid tumor, primary ascites cells, non-humanized PDX solid tumor and non-humanized PDX ascites cells for select patients. FFPE sections of matched patient solid tumor was obtained through the UNM Human Tissue Repository. For patient solid tumor samples, RNA isolation was performed by the UNMCCC Analytical and Translational Genomics (ATG) Shared Resource, as described previously^20, 21^. Briefly, total RNA was isolated from slide-mounted FFPE sections using the RNeasy FFPE kit (Qiagen, Hilden, Germany). For PDX samples, snap frozen solid tumor was incubated in RNAlater™-ICE Frozen Tissue Transition Solution (Invitrogen) for 24 hours at -20°C prior to DNA/RNA extraction with AllPrep DNA/RNA Mini Kit (Qiagen). Cryopreserved patient ascites cells and PDX cancer spheroids were thawed and rinsed in serum free RPMI before RNA/DNA extraction. cDNA synthesis and library preparation were performed in the UNMCCC ATG Shared Resource using the SMARTer Universal Low Input RNA Kit for Sequencing (Clontech, San Jose, CA) and the Ion Plus Fragment Library Kit (Life Technologies) as previously described^21^.

### RNA-seq Analysis

Low quality and non-human RNA-seq reads were identified and removed from the analysis pipeline using the Kraken suite of quality control tools^22, 23^. High-quality, trimmed, human RNA-seq reads were aligned to the human genome (GRCh37; hg19) using TMAP (v5.0.7) and gene counts were calculated using HT-Seq as previously described^21^. Gene Set Enrichment Analysis (GSEA) comparing PDX samples, primary OvCa ascites, and primary OvCa solid tumors was analyzed using the GSEA software (http://www.gsea-msigdb.org/gsea/index.jsp)^24^. 1,742 genes in the OvCa dataset were compared to 7,871 gene sets from the Molecular Signature Database after filtering out for gene set size (minimum 15, maximum 500 genes/set). For comparison of PDX and primary ovarian cancer gene expression, nine solid tumor or ascites PDX samples were compared to thirteen primary or omental ovarian tumors, and ten primary ascites samples. Samples were divided into three groups: PDX, primary ascites or primary solid tumor. Genes with significantly altered expression between groups were tabulated.

### Humanized NBSGW mice

Cryopreserved human cord blood-derived CD34+ cells (STEMCELL Technologies, Cambridge, MA) were rapidly thawed at 37°C, resuspended in media (RPMI +1% human serum albumin), centrifuged (300 x g, 10 min. at RT), and rinsed in media. After centrifugation, the cell pellet was resuspended in 1 ml of Stemline® II Hematopoietic Stem Cell Expansion Medium (Sigma-Aldrich), supplemented with 0.1 μg/ml Human Recombinant SCF (STEMCELL). Cells were incubated overnight in a standard CO_2_ incubator at 5% CO_2_. Prior to engraftment, cells were centrifuged (300 x g for 10 min. at RT) and resuspended in PBS. 2.5 × 10^5^ CD34+ cells were administered by retro-orbital injection into 3-4 week-old female NBSGW mice. Pooled donor samples were used to limit between-donor variability. Peripheral blood was drawn from humanized NBSGW mice at 8 weeks post-engraftment to characterize humanization (% human CD45+ cells/ total mouse CD45+ cells + human CD45+ cells) and human immune cell subpopulations by flow cytometry (**Supplementary Fig S1A**). Antibodies are listed in **Supplementary Table S1**. Mice with greater than 25% human CD45+ cells in peripheral blood were considered optimally humanized. Percent humanization at 8 weeks post-engraftment ranged from 40% to 84% (**Supplementary Fig S1B**). Mice were divided into three groups for engraftment of three PDX models. Humanization was not significantly different between the groups.

### Humanized PDX mice

Humanized NBSGW mice were injected 10 weeks post-CD34+ cell engraftment with fresh ovarian cancer spheroids from three different PDX models (PDX3 passage 2 (P2), PDX9 P4, and PDX18 P6). 5 × 10^6^ cells/mouse were injected IP to seed orthotopic disseminated ovarian cancer. Mice were weighed weekly and monitored for wasting or abdominal distention. Once mice showed signs of disease, they were monitored daily and sacrificed at a humane endpoint. Peripheral blood was collected at two and six weeks post-tumor engraftment and blood and ascites fluid were collected at endpoint. All tumors were removed and the mass of the total tumor burden was recorded.

### IHC of humanized solid tumors

Solid tumors and spleens from PDX models (n=2-3 per group) were formalin fixed at necropsy. HuPDX spleens were used for IHC optimization of anti-human antibodies in mouse tissue. Fixed tissue was paraffin embedded and sectioned by the UNM HTR and immunohistochemistry was performed on the Ventana Discovery Ultra Platform using immunoperoxidase labeling. Serial sections were processed for each marker. Antibodies are listed in **Supplementary Table S1**. For digital pathology and HALO analysis (Indica Labs, Albuquerque, NM), quantitative analysis algorithms were optimized to detect intratumoral infiltrating human immune cells. Immune cells were classified as intratumoral or extratumoral based on a tumor tissue mask based on labeling with a human mitochondrial marker or H&E staining. After omitting areas with obvious staining artifacts, all cells within the tumor were identified based on the nuclear hematoxylin counterstain and positive cells were tallied as weak, medium or strong staining. The average number of positive cells/100,000 cells were plotted for each group. For analysis of tumor vascularization, tumor vessels were labeled with anti-mouse CD31 antibody and slides were digitally scanned. The vessel area was determined using the tissue classifier analysis software on the HALO platform. Vascular density was reported as percent vascular area and was calculated based on analysis of whole sections from huPDX and non-huPDX tumors.

### Cytokine analysis of ascites fluid and peripheral blood plasma

For cytokine array analysis, frozen cell-free ascites fluid and peripheral blood plasma samples were submitted to Eve Technologies (Calgary, Canada) for analysis on their Human Cytokine/Chemokine Array 48-Plex. Forty-eight cytokines, chemokines, and growth factors were analyzed in duplicate from 100 μl of sample. Human M-CSF and VEGF-A levels were measured in patient and PDX cell-free ascites fluid using the Human M-CSF ELISA Kit or Human VEGF-A ELISA kit respectively (RayBiotech, Peachtree Corners, GA). HuPDX cytokine levels were compared to published LINCOplex microarray data from paired ascites and plasma samples from ovarian cancer^25^. Macrophage-derived chemokine (MDC) values were compared to published data from 93 ovarian cancer patients^26^. For uniformity, average and SEM for MDC levels were estimated from the listed median and interquartile range using an online estimator^27^.

### Statistical Analyses

Statistical methods were applied to the cytokine array data to determine whether observed cytokine measurements differed according to two primary factors: ascites vs. plasma sample source and huPDX vs. non-huPDX. First, a linear mixed effects model was fit to the data, treating specific cytokines as a repeated measure factor. This made it possible to test the global significance of cytokine-specific differences among the various grouping factors. It also enabled the estimate of average between-group differences across all cytokines. After identifying factors where there was evidence of statistical differences between groups, we fit a separate analysis of variance model to each cytokine to test the significance of the between-group differences and estimate the within-group means for each cytokine. For survival curve comparisons, analyses were performed using GraphPad Prism software and used a Log-rank Mantel-Cox test to determine significant differences between huPDX and non-huPDX.

## RESULTS

### Ovarian cancer PDX in immunocompromised mice lack immune cells that are crucial for establishing a supportive tumor microenvironment in solid ovarian tumors and ascites fluid

To establish ovarian cancer PDX models that sample the genetic heterogeneity of the UNMCCC patient population, cancer spheroids isolated from the malignant ascites of confirmed high-grade serous ovarian cancer patients were engrafted into the peritoneal cavity of NSG mice. PDX developed from 6 out of 9 patient samples for a 67% take rate, similar to rates reported in the literature for solid ovarian cancer PDX (68%-74%) and higher than reported rates for spheroid derived PDX orthotopic tumors (31%)^3, 4, 6, 28^. Our PDX models developed tumors within the omentum and mesentery, near the ovaries, and in the perigonadal fat pads with occasional spread to the liver and diaphragm. The dissemination throughout the peritoneal cavity is similar to that seen in ovarian cancer patients.

In a comparison of RNA-seq gene expression data from primary ovarian cancer and PDX samples, 43% of down-regulated genes in PDX samples were either known macrophage markers (e.g.CD68, CD163), involved in macrophage polarization (e.g. MALAT1, NEAT1, IL10RA) or responsible for immune infiltration (e.g. THBS1, THBS2, ITGA5, and collagens) (**Figure 1A**). Furthermore, RNA-seq analyses of unmatched ovarian cancer primary ovarian tumors and primary malignant ascites revealed an IFNγ-stimulated inflammatory macrophage signature shared across ascites samples but not in solid tumor samples (**Figure 1B,C**). This supports the existence of two immune microenvironments, one in solid tumors and one in the ascites fluid. These distinct immune environments, which are absent in conventional PDX models, may influence ovarian cancer growth, dissemination, and response to therapies.

**Figure 1.**
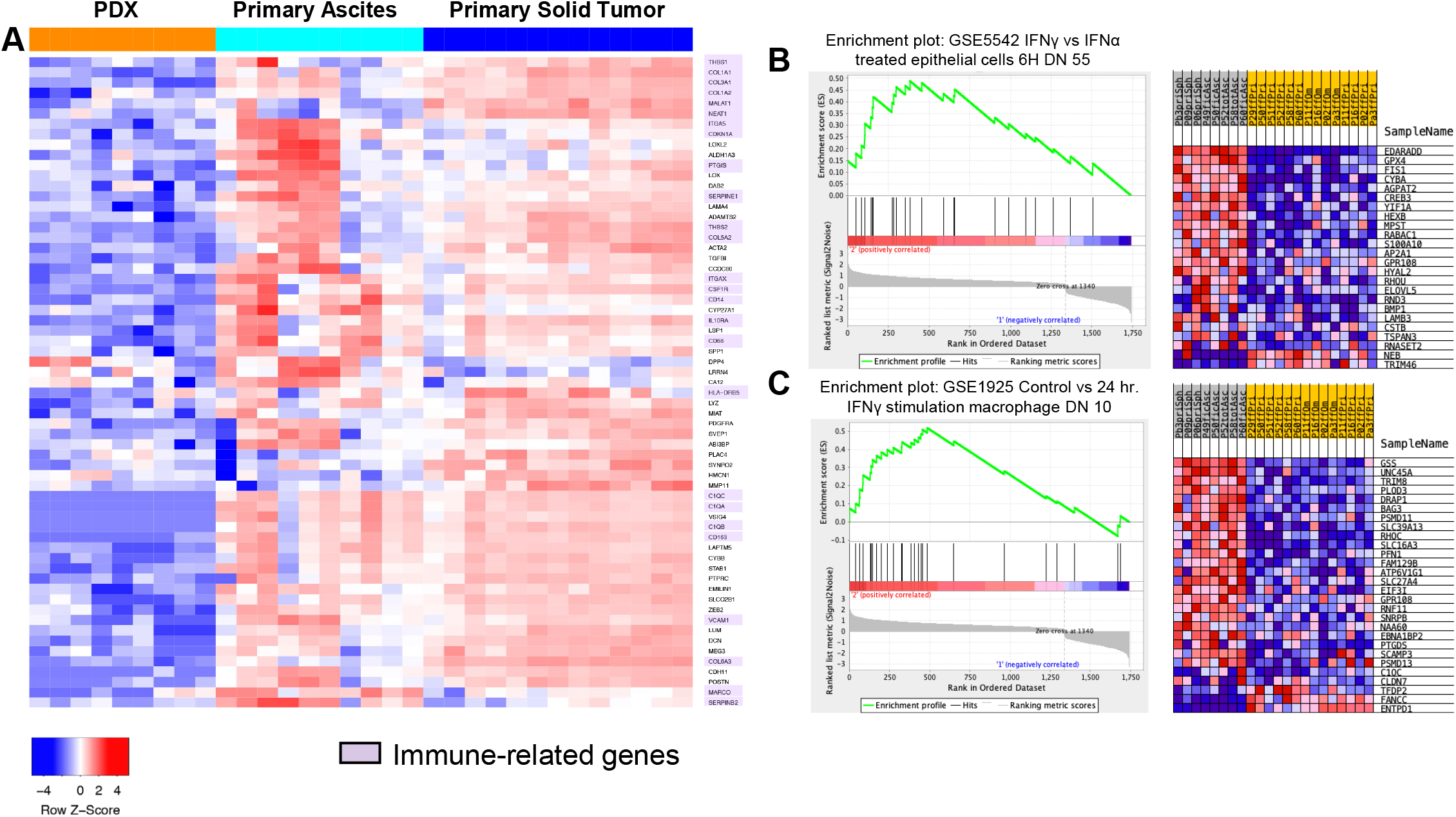
Analysis of ovarian cancer patient samples reveals an inflammatory signature in malignant ascites samples. **A**. Of the differentially expressed genes that show significantly lower expression in non-humanized PDX samples (orange) compared to primary OvCa ascites cells and primary OvCa solid tumor samples, 43% are immune-related genes (highlighted in purple). **B-C**. Gene Set Enrichment Analysis (GSEA) comparing primary ascites (n=8) and primary solid tumors (n=14) shows enrichment in genes associated with IFNγ stimulation in primary ascites.

### Establishing Ovarian Cancer PDX models with a reconstituted humanized immune system

To examine the influence of the tumor immune microenvironment on PDX disease progression, we chose three PDX models that demonstrated robust growth in immunocompromised mice, two platinum-sensitive PDXs (PDX3 and PDX18) and one platinum resistant PDX (PDX9) to engraft in huNBSGW mice. Characteristics of the three patients are provided (**Supplementary Table S2**). These PDX were injected into the peritoneal cavities of humanized NBSGW mice engrafted with human cord blood-derived CD34+ HSCs from the same donor pool to establish a uniform immune background. Non-humanized NBSGW injected with the same three PDX served as non-humanized PDX controls. The huPDX study design is diagrammed in **Figure 2A**.

**Figure 2.**
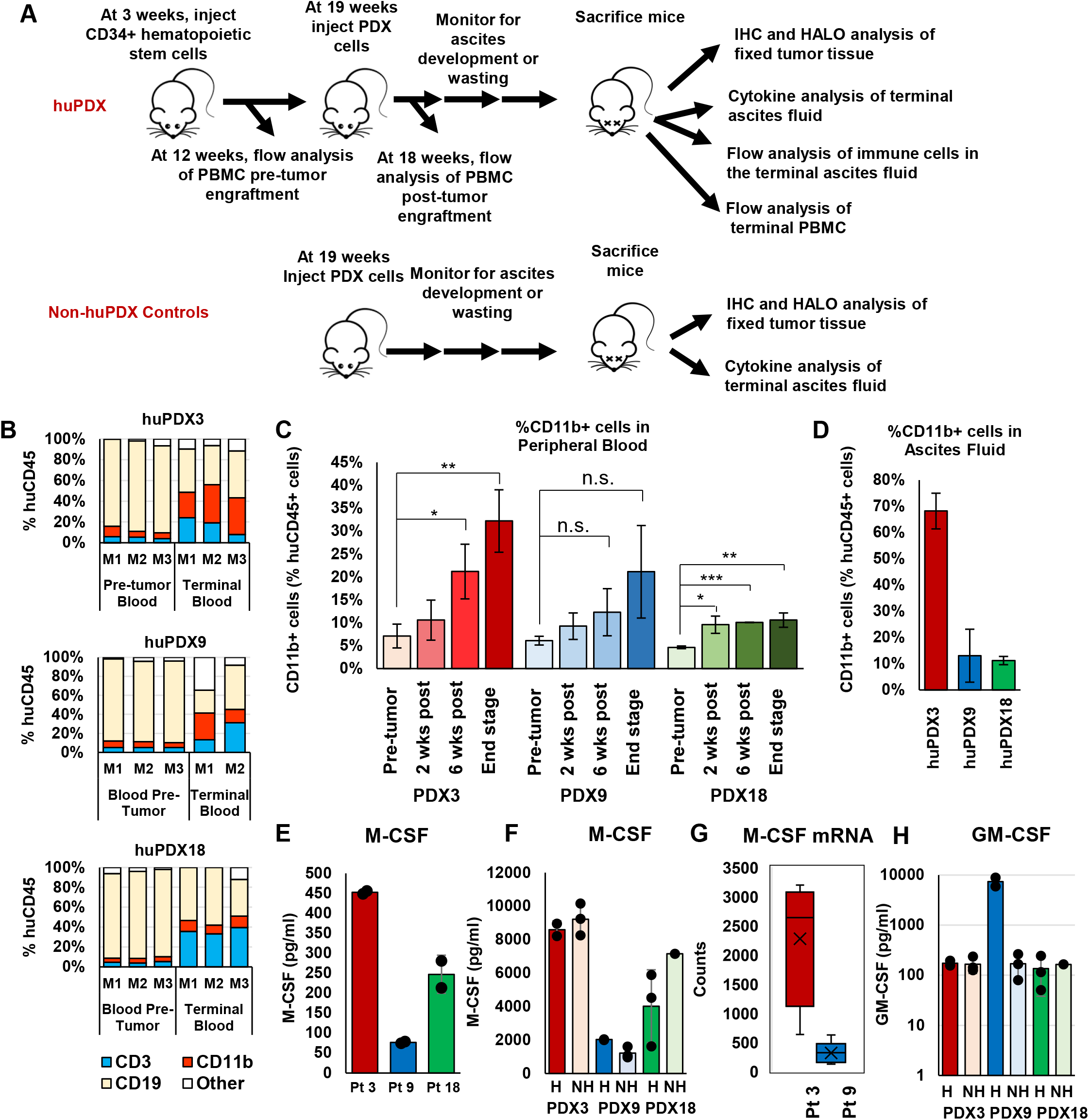
High M-CSF production influences the development of myeloid cells in humanized NBSGW PDX. **A**. Diagram of the huPDX study. **B**. Change in human CD45+ cell populations in peripheral blood pretumor engraftment versus end stage for each mouse. CD19+ B cells, CD3+ T cells, and CD11b+ myeloid cells make up the majority of human immune cells in the blood. **C**. Average percentage of human myeloid cells in the peripheral blood of huPDX pre- and post-tumor challenge (values are the average +/-STDEV. p-values are based on an unpaired two-tailed Student’s t-test. ***p<0.0005, ** p<0.005, *p<0.05, n.s. not significant). **D**. Percentage of CD11b+ human myeloid cells in huPDX ascites fluid at end stage. **E**. M-CSF gene expression levels in patient samples as determined by RNA-seq analysis of primary patient samples and non-huPDX samples. **F**. M-CSF concentration in patient ascites fluid measured by ELISA. Samples were assayed in duplicate. Error bars are standard deviation. **G**. M-CSF concentrations and **H**. GM-CSF concentrations in PDX ascites fluid measured by cytokine array. Dots represent the value for each sample. Bars are the average level and error bars are SEM. H=humanized NH= non-humanized

### Engrafted ovarian cancer PDX models drive increased myeloid cell differentiation in humanized NBSGW mice

Testing the impact of PDX tumor challenge on the development of engrafted human leukocytes revealed a striking change in the human CD45+ cell subpopulations in the peripheral blood of huNBSGW (**Figure 2B-C**). Notably, tumor challenge led to a significant increase in the percentage of human myeloid cells as highlighted in **Figure 2C**. Before tumor challenge, CD11b+ myeloid cells constituted on average 6% +/-1.8% of the human CD45+ cells in huNBSGW peripheral blood. At end stage, the average percentage of myeloid cells had risen to 21.3% +/-10.8%. Myeloid cells also made up a large proportion of human leukocytes within the ascites fluid of huPDX (**Figure 2D**). HuPDX3 had the highest percentage of human myeloid cells accounting for 32% +/-6.8% of the human CD45+ cells in the peripheral blood and 68.2% +/-11.3% in the ascites fluid. The increase in myeloid cells seen in PDX3 was confirmed with another donor pool of CD34+ cells (**Supplementary Figure S1C**) demonstrating that the increase in myeloid cells was not donor dependent, but instead driven by the tumor cells. Although CD3+ T cells increased over time, with a concomitant decrease in human CD19+ B cells (**Figure 2B)**, non-tumor bearing humanized NBSGW showed similar changes over time (**Supplementary S1D**). Others have reported similar findings in humanized models, indicating that the effect is not human tumor dependent but a characteristic of humanized models^29^. Thus, growth of ovarian cancer PDX tumors in huNBSGW does not affect T and B cell differentiation, but improves myeloid differentiation and results in the accumulation of myeloid cells in the peripheral blood and ascites fluid.

### Engrafted ovarian cancer PDX models have elevated levels of human cytokines in the ascites fluid

To characterize human cytokine profiles of huPDX models, end stage acellular ascites fluid and blood plasma from huPDX and non-huPDX were analyzed on a 48-plex human cytokine/chemokine array. Plasma from a non-tumor bearing huNBSGW mouse served as a control for the production of human cytokines by human immune cells in the absence of tumor cells. The control huNBSGW had high plasma levels (>100 pg/ml) of human IFNα2, IL12p40, MDC, MIG, and RANTES and lower but detectable levels of ten other cytokines (**Supplementary Figure S2**). Plasma from a non-humanized, non-tumor bearing NBSGW served as a negative control to ensure that mouse cytokines were not detected by the human array. Thirty-five human cytokines had significantly higher levels in the PDX ascites fluid samples relative to plasma, demonstrating that cytokines produced in the ovarian cancer tumors are concentrated in the peritoneal microenvironment (**Supplementary Table S3**). Overall, cytokines were 2.42 log-concentration units higher in ascites vs. plasma samples (S.E.=0.30, p<0.001). This agrees with a previous report finding higher concentration of cytokines in the ascites fluid versus plasma of ovarian cancer patients^25^.

### PDX tumor cells produce human M-CSF and GM-CSF that can influence myeloid differentiation

The observed increase in myeloid cells suggested that human cytokines produced by the ovarian cancer cells influence human immune cell differentiation within huPDX models. Humanized mice do not normally have robust myeloid cell differentiation due to the lack of cross reactivity of mouse myeloid differentiation factors, mouse M-CSF and mouse GM-CSF, with human receptors. Analyses of M-CSF cytokine levels in primary patient and PDX ascites detected high levels of human M-CSF in all samples (**Figure 2E**,**F**). There were no significant differences in M-CSF levels comparing tumor bearing huPDX and non-huPDX models (**Figure 2G**), confirming that human M-CSF is produced by ovarian cancer cells. M-CSF secretion levels are correlated with transcription levels based on RNA-seq data analyses comparing the PDX models with the highest and lowest levels of human M-CSF, PDX3 and PDX9 respectively (**Figure 2E**). GM-CSF was also detected in the ascites fluid of all PDX mice (**Figure 2H**). The data shows that ovarian cancer huPDX models produce myeloid differentiation factors that improve human myeloid reconstitution beyond what has been characterized in non-tumor-bearing huNBSGW.

### HuPDX models recapitulate the cytokine milieu of ovarian cancer patient ascites

A large number of cytokines have been reported as elevated in ovarian cancer ascites samples including IL-6, IL-8, IL-10, IL-15, IP-10/CXCL10, MCP-1/CCL2, Mip1α/CCL3, Mip1β/CCL4, MDC and VEGF^25, 26, 30^. All of these cytokines were found at high levels in the huPDX ascites samples (**Table 1)**, including chemokines that recruit immune cells to the tumor microenvironment such as MCP-1/CCL2 that recruits monocytes^31^, MDC that recruits regulatory T cells^26^, and MIG1 (CXCL9) and IP-10 (CXCL10) that recruit effector memory T cells^32, 33^. Importantly, IL-10, an important immunosuppressive cytokine, is absent from the ascites fluid of non-huPDX models, but is detectable in all three huPDX models. Thus, the huPDX immune environment recapitulates that found in human ovarian cancer patients.

**Table 1.** Cytokines elevated in the ascites fluid of ovarian cancer patients are also present in the huPDX ascites fluid. Nearly all cytokines were higher in huPDX models compared to non-huPDX. Values are expressed as average concentrations in pg/ml. All ovarian cancer ascites fluid values are from Giuntoli et al., 2009 except MDC that is an estimate of the average from Wertel et al, 2015. Color reflects the relative concentration levels with magenta higher and blue lower.

|  | OvCa Ascites Fluid<br>Published Study |  | huPDX3 |  | huPDX9 |  | huPDX18 |  |
| --- | --- | --- | --- | --- | --- | --- | --- | --- |
|  | Average<br>(pg/ml) | SEM | Average<br>(pg/ml) | SEM | Average<br>(pg/ml) | SEM | Average<br>(pg/ml) | SEM |
| IL-6 | 14,845 | 4,658 | 1,682 | 1,274 | 13,457 | 2,499 | 175 | 64 |
| IP-10 | 11,689 | 1,645 | 165,153 | 29,231 | 14,582 | 10,295 | 10,831 | 8,860 |
| MCP-1 | 1,456 | 365 | 38,958 | 3,038 | 19,604 | 9,961 | 3,589 | 3,174 |
| IL-8 | 1,122 | 309 | 7,677 | 1,143 | 16,958 | 10,763 | 1,268 | 419 |
| Mip1β | 730 | 184 | 230 | 98 | 839 | 0 | 27 | 7 |
| MDC | 435 | 22 | 940 | 244 | 2,292 | 181 | 576 | 105 |
| IL-10 | 138 | 25 | 88 | 60 | 17,177 | 1,352 | 89 | 19 |
| Mip1α | 60 | 7 | 149 | 52 | 6,393 | 4,089 | 50 | 7 |
| IL-15 | 42 | 10 | 209 | 39 | 163 | 23 | 207 | 60 |

### Comparison of huPDX versus non-huPDX models identifies multiple origins of human cytokine production

In the huPDX models, human cytokines can be produced by the cancer cells or by the engrafted human immune cells. Comparison of huPDX and non-huPDX revealed three types of cytokine expression patterns: PDX-intrinsic cytokines, immune cell-dependent cytokines, and immune cell-influenced cytokines. Examples of all three types are shown in **Figure 3**. PDX-intrinsic cytokines are produced by the cancer cells and were present in non-huPDX and huPDX ascites fluid (**Figure 3A**). Twenty PDX-intrinsic cytokines were identified in all three PDX models (**Figure 3D**). Other cytokines had significantly higher levels in the huPDX vs non-huPDX samples. Statistical analyses comparing the least squares means of cytokine levels in huPDX vs non-huPDX ascites samples demonstrated a significant increase in the levels of 28 cytokines in huPDX ascites. These cytokines were subdivided into immune cell-dependent factors that were only present in the huPDX samples (**Figure 3B**) and immune cell-influenced factors that had lower levels in the non-huPDX with significantly higher levels in the huPDX (**Figure 3C**). Cytokines detected in the PDX ascites fluid are listed in **Figure 3 D-F**. A large number of cytokines are PDX-intrinsic highlighting the potential for cancer cells to influence human immune cells within the tumor microenvironment. Human immune cells produce important cytokines such as IL-10 and increase the concentration of many cytokines within the ascites fluid emphasizing the impact of human immune cells on the tumor microenvironment.

**Figure 3.**
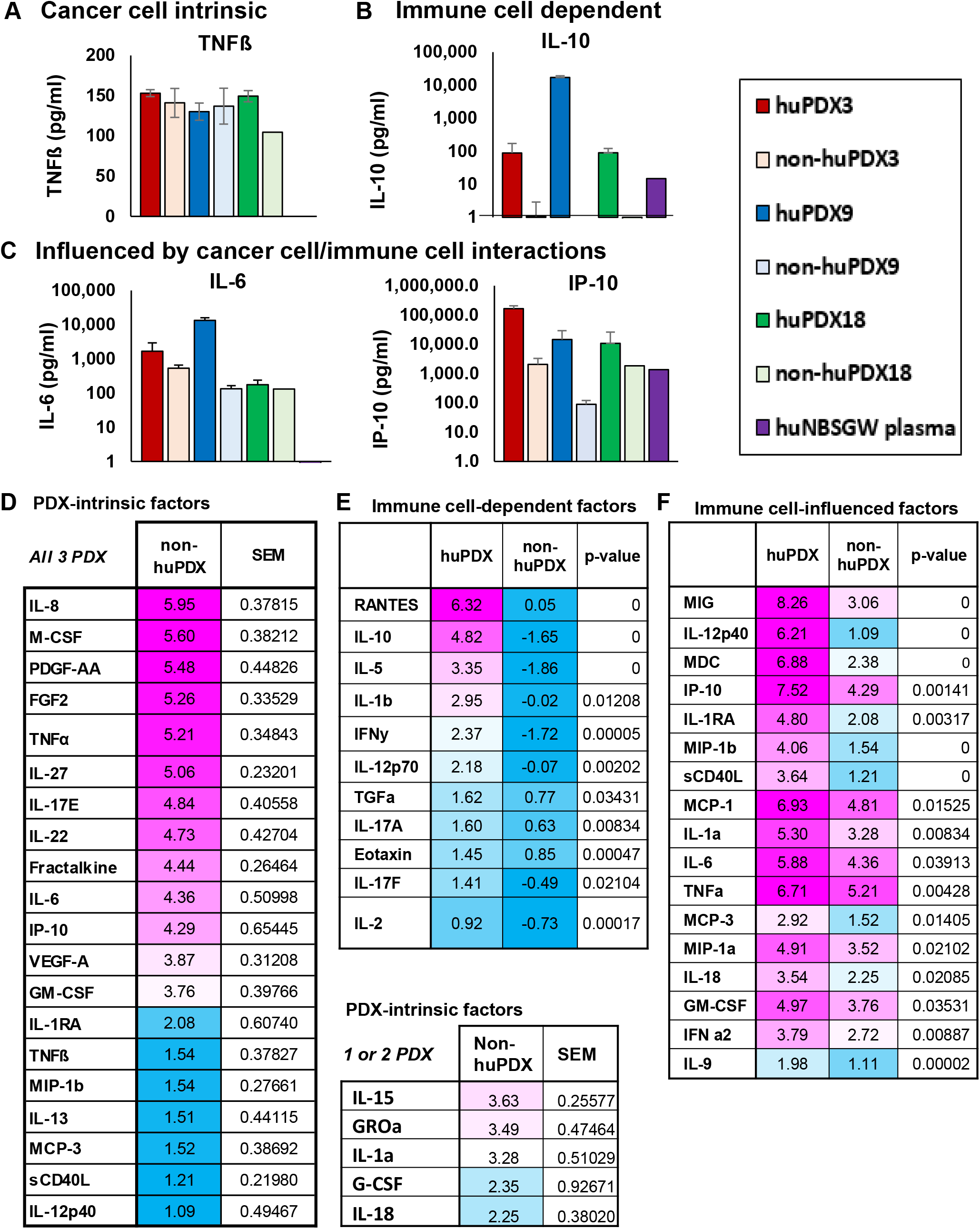
Analysis of human cytokine levels in huPDX and non-huPDX models indicates their origin. A-C. Examples of the subtypes of cytokines in huPDX. **A**. PDX-intrinsic cytokines. **B**. Immune cell-dependent cytokines. **C**. Immune cell-influenced cytokines. **D-F**. Tables of cytokine levels in ascites fluid. Levels are presented as the natural log of the Least Squares Means. **D**. PDX-intrinsic cytokines (levels >1 in non-huPDX). Factors with detectable levels in only 1 or 2 PDX models are listed separately. **E**. Immune cell-dependent cytokines with non-huPDX levels <1. **F**. Factors with detectable levels in the non-huPDX, but significantly increased levels in the huPDX.

### Human tumor-associated macrophages (TAMs) and tumor infiltrating leukocytes (TILs) are detected in huPDX tumors

Tumor infiltrating immune cells can influence disease progression, patient prognosis, and response to therapies. To determine whether human immune cells were recruited to solid tumors, formalin fixed paraffin embedded tumors from huPDX and non-huPDX mice were serially sectioned and labeled using human-specific immune cell markers to identify CD68+ TAMs (**Figure 4**) and CD3+ TILs (**Figure 5**). Quantification of labeled immune cells was performed using HALO software (**Supplementary Fig S3A**). Non-huPDX tumor sections were used as negative controls **(Supplementary Fig S4)**. Both TAMs and TILs were identified in tumors from all three PDX models. In PDX3, TAMs had infiltrated into cancer cell clusters throughout the tumors, while in PDX9 and PDX18 TAMs were primarily localized to stromal tissue surrounding the cancer cell clusters. Quantification of CD4+ vs CD8+ TILs revealed PDX-specific differences in TIL infiltration (**Figure 5**). HuPDX3 TILs were primarily CD4+ (CD4+/CD8 average ratio of 4.7 +/-3.3), which correlates with poor survival in ovarian cancer patients^34^. HuPDX18 had similar levels of CD4+ and CD8+ T cells (CD4+/CD8+ average ratio of 0.8 +/-0.6). HuPDX9 TIL counts were minimal with counts comparable to the non-huPDX tumor controls. Though all three huPDX models had detectable TAMs and TILs, the density and localization of tumor-infiltrating immune cells appears to be PDX-dependent.

**Figure 4.**
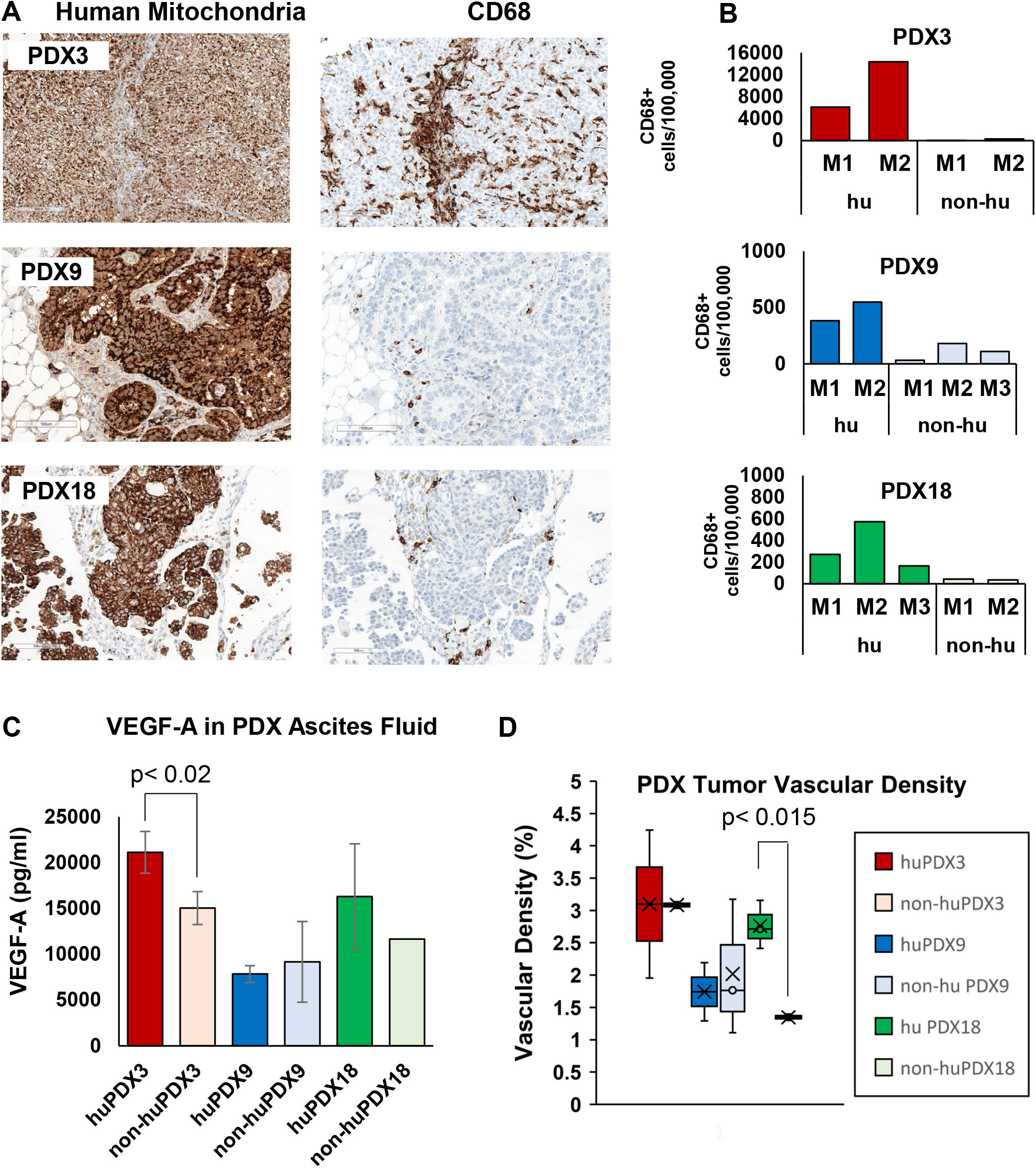
Detection of TAMs in huNBSGW PDX tumor sections. **A**. Serial sections of huNBSGW PDX tumor tissue were labeled with human-specific antibodies against a mitochondrial protein to label all human cells and the TAM marker CD68. B. Quantification of the number of TAMs using HALO software to identify CD68+ cells. Values are # positive cells/100,000 cells. **C**. Concentration of VEGF-A in PDX ascites fluid was measured by ELISA. **D**. The quantification of vascular density (% vascular area) across whole sections of huPDX and non-huPDX samples. Values are the average vascular densities among samples (n=2-3). huPDX18 sections showed increased vascular density compared to non-huPDX18 (p<0.015, unpaired t-test).

**Figure 5.**
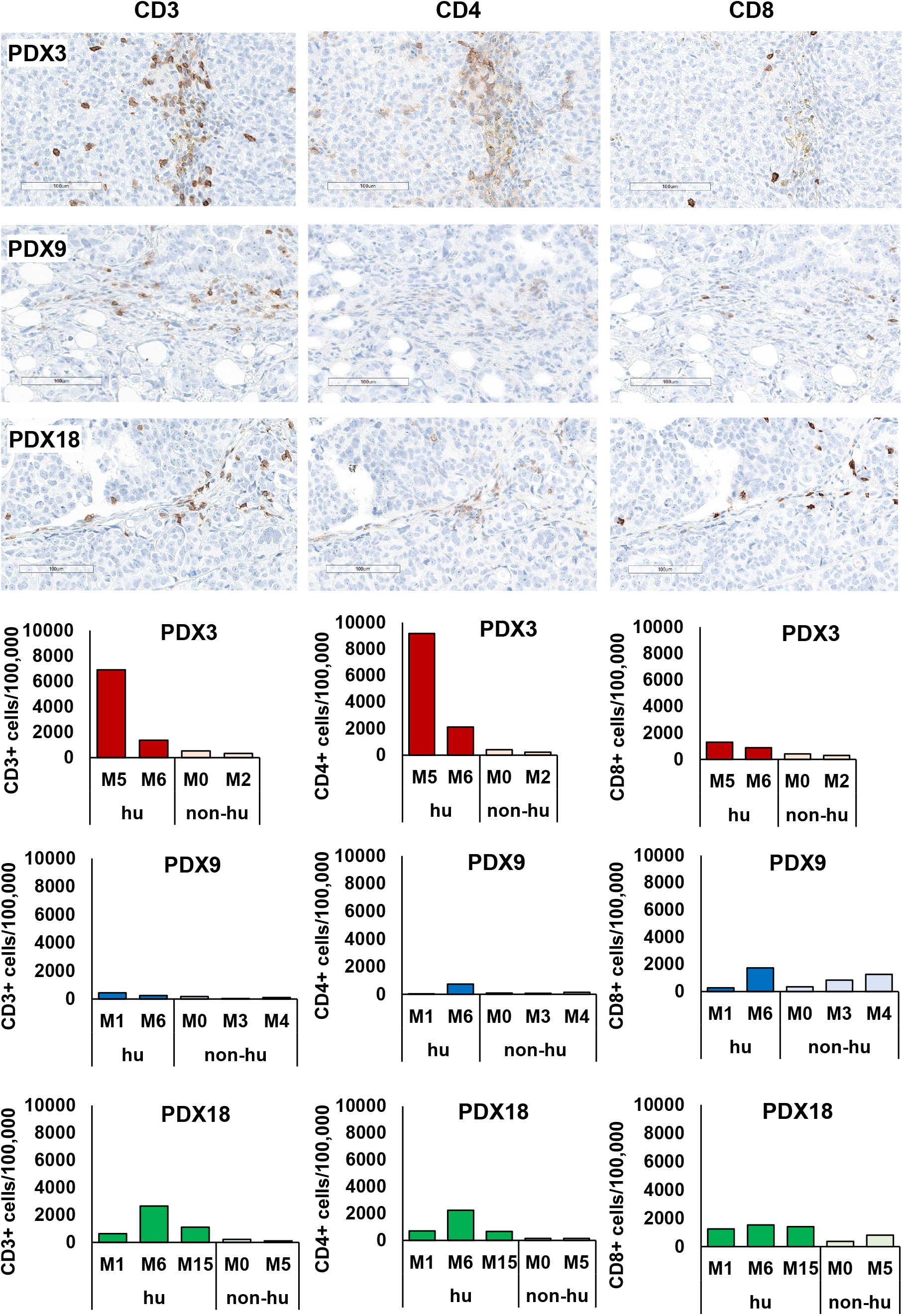
Human TILs in PDX tumors. **A**. Tumor sections from huPDX were labeled for human T cell markers CD3, CD4, and CD8. Shown are scanned images of labeled sections at 20X magnification. Representative images are shown for huPDX3 and huPDX18. For huPDX9, the image shown is a rare group of TILs in a mostly negative section. **B**. Quantification of the number of TAMs using HALO software. Numbers are given as # positive cells/100,000 cells.

It was previously reported that the presence of TAMs can increase tumor vascularization by boosting VEGFA production^35^. Indeed, VEGF-A levels were significantly higher in the ascites of huPDX3 compared to non-huPDX3 and huPDX18 levels also trended higher (**Figure 4C**). To examine whether tumor vascularization increased in the huPDX models, tumor sections were labeled with a CD31 antibody and vascular density was quantified (**Figure 4D**). PDX18 showed a significant increase in vascular density with humanization, suggesting that local VEGF production by human immune cells may have boosted VEGF levels enough to impact vascularization in this model. No change in vascular density was observed in PDX3 or PDX9. Therefore, further studies are needed to affirm a primary role for VEGF secretion by TAMs and tease out other factors that may be involved.

Analysis of huPDX tumors reveals that human immune cells can be recruited by PDX cells to recapitulate the tumor immune microenvironment in huPDX. Comparison of three huPDX models shows that huPDX can represent between-patient variability in immune cell recruitment and tumor vascularization.

### Engrafted human immune cells influence PDX disease progression

Similar to what is observed with the non-huPDX; survival times for the huPDX models were patient-specific. PDX that consistently demonstrated shorter survival times (PDX3 and PDX9), also showed more aggressive disease in huPDX. However, for all PDX models, huPDX mice had shorter survival times than the non-huPDX derived from the same patient (**Figure 6A**), indicating that the tumor-promoting role of human immune cells is dominant over the anti-tumor role in huPDX models. Tumors did not simply grow faster in huPDX mice, since the total tumor mass measured at endpoint was lower in huPDX compared to non-huPDX (**Figure 6B**). Humanized PDX3 showed signs of anemia at end stage, likely due to enhanced myeloid differentiation^13^, but all huPDX3 mice presented with malignant ascites prior to sacrifice indicating disease progression. Other disease characteristics with relevance to human disease contributed to early sacrifice in the huPDX mice. For instance, two of three huPDX9 mice presented with bowel obstruction prior to ascites development, while none of the non-huPDX9 mice showed evidence of bowel obstruction. All non-huPDX9 were sacrificed due to abdominal distension with ascites volumes similar to the other PDX groups (**Figure 6C**). Since malignant bowel obstruction is a common complication of ovarian cancer affecting up to 51% of patients with recurrent disease, future studies with larger cohorts using huPDX9 will provide a model for studying the underlying causes of bowel obstruction in patients^36^. HuPDX18 accumulated malignant ascites much earlier than the non-huPDX18 requiring early sacrifice. Therefore, the influence of immune cells on ascites accumulation will be studied using the huPDX18 model. Reducing ascites accumulation is particularly important for ovarian cancer palliative care. The presence of human immune cells clearly influences disease progression in complex and PDX-specific ways. Collectively, the use of huPDX models affords new opportunities to examine disease relevant complications that may be driven by specific tumor cell characteristics.

**Figure 6.**
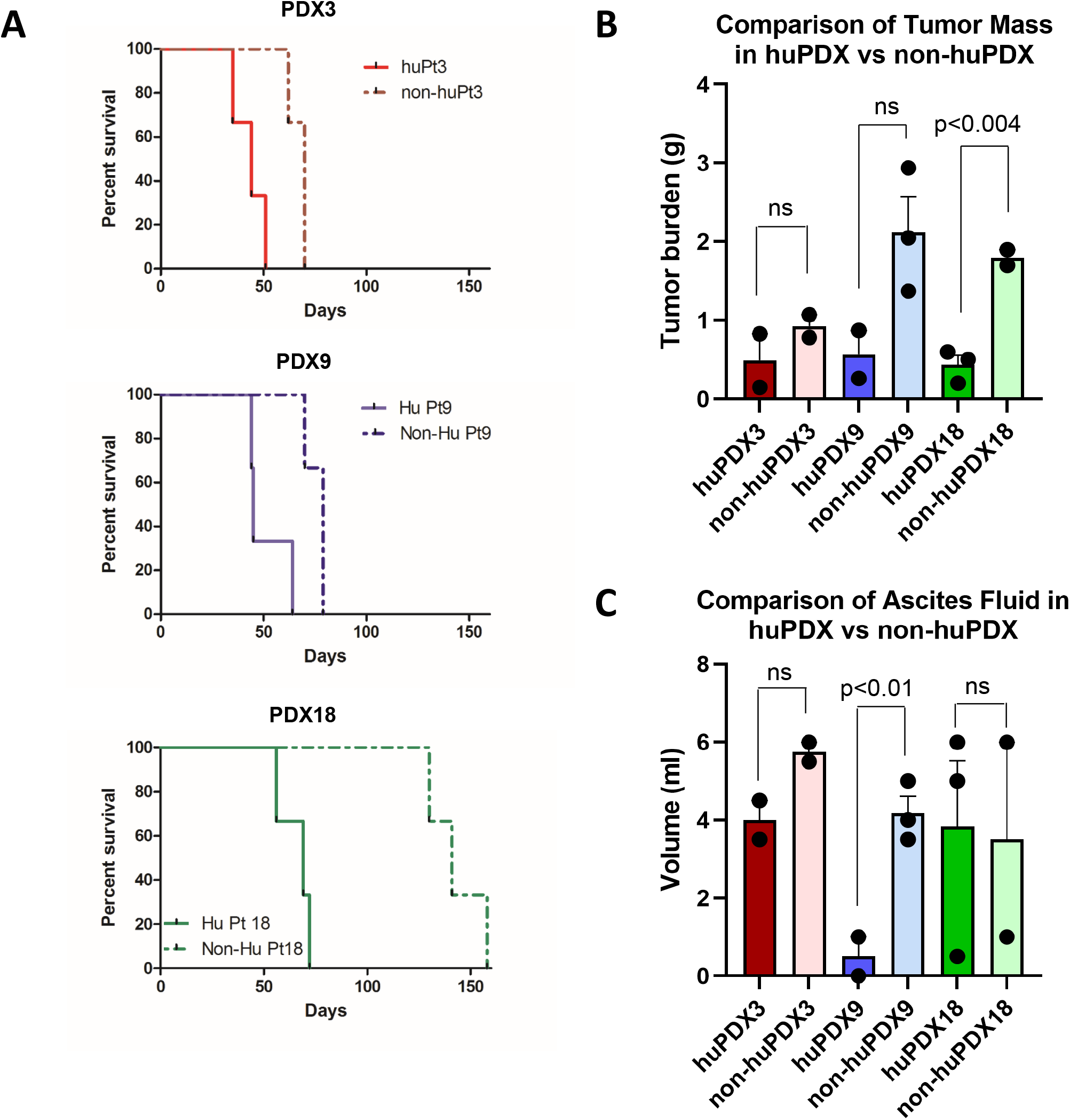
Survival time is reduced in huNBSGW PDX models. **A**. huNBSGW PDX 3, 9 and 18 were engrafted with the same pooled donor CD34+ cells. Survival time was PDX-dependent with PDX3 and 9 having a shorter survival time compared to PDX18. Survival curves are significantly different between huPDX and non-huPDX for all three PDX models (p<0.025, Log-rank test). **B**. Solid tumor mass (in grams) of huPDX and non-huPDX at necropsy. Error bars are SEM. **C**. Ascites fluid volume at necropsy. The ascites fluid volume was higher in non-huPDX for PDX3 and PDX9, but not PDX18.

## DISCUSSION

In ovarian cancer, the importance of TAMs and MDSCs in disease progression, metastasis and therapeutic response has been well established^8, 9, 37^. Yet current patient-derived models for testing therapeutic response, CDX and PDX in immunocompromised mice, lack a tumor immune microenvironment. Thus, preclinical testing can miss any immunomodulatory effects of novel therapies. To better reflect the immune microenvironment, recent studies have considered humanized mouse models. However, standard humanized NSG mice have demonstrated minimal differentiation of human myeloid populations after human HSC engraftment^12^, raising the question of whether even humanized mice can recapitulate the tumor immune microenvironment. Here, we detail the utility of a humanized mouse model of orthotopic ovarian cancer using the NBSGW mouse strain that obviates the need for pre-engraftment irradiation, while improving engraftment and differentiation of human myeloid cells^18^. This is the first study reporting the use of humanized NBSGW mice for cancer research and the first to characterize a humanized immune system (HIS) mouse model of ovarian cancer.

We show that ovarian cancer huPDX enhance human myeloid cell differentiation through the production of human cytokines such as M-CSF and GM-CSF by ovarian cancer cells. Therefore, our model reconstitutes myeloid cells without requiring constitutive expression of human cytokines by the mouse host. Apart from factors involved in differentiation and recruitment of macrophages, other human immune differentiation factors were detected in huPDX models, such as G-CSF that is important for the activation of neutrophils and IL-15 that is responsible for the differentiation and activation of human NK cells. This suggests that our huPDX models may support a broader repertoire of human immune cells. Furthermore, we show PDX-specific recruitment of human CD68+ TAMs and CD3+ TILS to huPDX solid tumors. The ability of huPDX to significantly increase the human macrophage population and recruit TAMs and TILs to solid tumors provides opportunities for identifying immune responses to therapies in a model that more broadly represents the complexity of the human tumor microenvironment.

By comparing non-huPDX and huPDX ascites fluid, we were able to determine which human cytokines were produced intrinsically by the cancer cells, and which required human immune cells. In particular, we identified PDX intrinsic production of human cytokines involved in the recruitment of macrophages to the tumor microenvironment. It is interesting to note that the PDX-intrinsic cytokines produced by ovarian cancer huPDX models are distinct from those produced by other cancer models producing high levels of IL-8, M-CSF, IP-10, MCP-1, VEGF-A, FGF-2 and PDGF-AA. A previous study using a 42-plex human cytokine array to analyze the serum of NSG mice engrafted with Nalm6-GFP leukemia cells reported high levels of PDX-intrinsic PDGF-AA and FLT-3L as well as lower levels of FGF-2, VEGF-A, and TNFα^38^. Tumors from humanized breast cancer models exhibited high levels of GM-CSF, IL-6, IL-8 and TNFα^39^. PDX-intrinsic cytokines may be specific to the tumor type.

Indeed, we find that cytokines previously identified in the ascites fluid of ovarian cancer patients are present at similar levels in huPDX ascites fluid, showing that these models recapitulate the human ovarian cancer peritoneal environment. Many of these huPDX-expressed cytokines, such as MCP-1, IL-6, and IL-10, can support an immunosuppressive peritoneal environment. MCP-1 (CCL2) is the main cytokine responsible for recruitment of CCR2+ immunosuppressive macrophages to tumors^31, 40, 41^. IL-6-dependent differentiation of CD4+ regulatory T cells has been stimulated by ovarian cancer ascites fluid *in vitro*^42^. Meanwhile, IL-10 downregulates T cell function and promotes immune tolerance^43^. High IL-10 levels (>24 pg/ml in ascites fluid), detected in all three huPDX models, are associated with significantly shorter progression free survival in ovarian cancer patients^44, 45^. These cytokines can also bind to receptors on the cancer cells and directly promote cancer invasion and metastasis^46-49^.

The effect of humanization on disease progression, with huPDX reaching endpoint faster than the non-humanized controls, suggests that, in the absence of treatment, the pro-tumor action of human immune cells dominates over anti-tumor immune responses. Previous work has noted that certain subcutaneous xenograft models grow faster in humanized mice, while others grow slower, indicating a heterogeneous immune response^7, 50^. Our finding that multiple ovarian cancer PDXs have shorter survival times in humanized mice suggests that immune effects on disease progression may be tumor type dependent. PDX-intrinsic cytokine production may explain the observed influence of the immune system on disease progression.

Comparison of three huPDX models demonstrated that the degree of immune cell recruitment is a characteristic of the individual PDX model and therefore dependent on patient genetic heterogeneity. PDX3 recruited larger numbers of TAMs, compared to PDX9 and PDX18. The localization of TAMs also varied among PDX models with TAMs in huPDX3 infiltrating into the epithelial tumor tissue, while TAMs in PDX9 and 18 remained within the stroma on the periphery of the tumors. PDX3 and PDX18 had higher numbers of T cells, while PDX9 had almost none. The high numbers of TILs in huPDX3 may be due to the high level of IP-10 (CXCL10). IP-10 has been reported to recruit TILs to ovarian cancer tumors^32^. Patient-specific differences in immune cell recruitment could be one of the factors influencing the variability in patient response to therapies. This highlights the need to include multiple PDXs in preclinical studies to model the differences in patient immune response.

Collectively, our data show that huPDX models in NBSGW mice reconstitute human myeloid cells and T-cells within the tumor microenvironment providing more clinically relevant models to examine cancer cell/immune cell interactions and test novel therapeutics. Our huPDX models are ideal for testing the response of genetically diverse ovarian cancer PDX to anti-cancer therapies including immunomodulatory agents such as immune checkpoint inhibitors and TAM targeting therapies.

## Authors’ Contributions

**Conception and design:** M.P. Steinkamp, I. Lagutina, A. Wandinger-Ness, L.G. Hudson, S.A. Ness, S.F. Adams

**Development of methodology:** M.P. Steinkamp, I. Lagutina, A. Wandinger-Ness

**Acquisition of data:** M.P. Steinkamp, I. Lagutina, F. Schultz, D. Burke,

**Analysis and interpretation of data:** M.P. Steinkamp, I. Lagutina, S.A. Ness, V.S, Pankratz, K.J. Brayer, F. Schultz

**Writing, review, and/or revision of the manuscript:** M.P. Steinkamp, A. Wandinger-Ness, L.G. Hudson, S.F. Adams, S.A. Ness, I. Lagutina

**Study supervision:** M.P. Steinkamp

## Acknowledgements

This research was supported by a Pilot Project Grant from the UNMCCC, a Pilot Project Grant from the UNM Pathology Department, and startup funds from the UNM Pathology Department and the UNMCCC. We gratefully acknowledge use of the UNMCCC Animal Models, ATG, Fluorescence Microscopy, Flow Cytometry, Biostatistics, and HTR Shared Resources as well as the NIH P30CA118100 grant that supports the UNMCCC and these shared resources. Additional technical assistance was provided by Russell Hunter, Shayna Lucero, Rachel Grattan, Jaimie Padilla (ATG), Wade M. Johnson (Flow), Cathy Martinez (HTR), and Monique Nysus (Animal Models).

## References

1. Weroha, S.J. et al. Tumorgrafts as <em>In Vivo</em> Surrogates for Women with Ovarian Cancer. Clinical Cancer Research 20, 1288–1297 (2014).

2. Scott, C., Becker, M.A., Haluska, P. & Samimi, G. Patient-derived xenograft models to improve targeted therapy in epithelial ovarian cancer treatment. Frontiers in Oncology 3 (2013).

3. Colombo, P.-E. et al. Ovarian carcinoma patient derived xenografts reproduce their tumor of origin and preserve an oligoclonal structure. Oncotarget 6, 28327–28340 (2015).

4. Liu, J.F. et al. Establishment of Patient-Derived Tumor Xenograft Models of Epithelial Ovarian Cancer for Preclinical Evaluation of Novel Therapeutics. Clinical cancer research : an official journal of the American Association for Cancer Research 23, 1263–1273 (2017).

5. Zayed, A.A., Mandrekar, S.J. & Haluska, P. Molecular and clinical implementations of ovarian cancer mouse avatar models. Chinese clinical oncology 4, 30–30 (2015).

6. Alkema, N.G. et al. Biobanking of patient and patient-derived xenograft ovarian tumour tissue: efficient preservation with low and high fetal calf serum based methods. Scientific reports 5, 14495 (2015).

7. Wang, M. et al. Humanized mice in studying efficacy and mechanisms of PD-1-targeted cancer immunotherapy. FASEB journal : official publication of the Federation of American Societies for Experimental Biology 32, 1537–1549 (2018).

8. Cheng, H., Wang, Z., Fu, L. & Xu, T. Macrophage Polarization in the Development and Progression of Ovarian Cancers: An Overview. Frontiers in Oncology 9 (2019).

9. Worzfeld, T. et al. The Unique Molecular and Cellular Microenvironment of Ovarian Cancer. Frontiers in Oncology 7 (2017).

10. Kipps, E., Tan, D.S.P. & Kaye, S.B. Meeting the challenge of ascites in ovarian cancer: new avenues for therapy and research. Nature reviews. Cancer 13, 273–282 (2013).

11. Billerbeck, E. et al. Development of human CD4+FoxP3+ regulatory T cells in human stem cell factor–, granulocyte-macrophage colony-stimulating factor–, and interleukin-3–expressing NOD-SCID IL2Rγnull humanized mice. Blood 117, 3076–3086 (2011).

12. Sippel, T.R., Radtke, S., Olsen, T.M., Kiem, H.-P. & Rongvaux, A. Human hematopoietic stem cell maintenance and myeloid cell development in next-generation humanized mouse models. Blood advances 3, 268–274 (2019).

13. Rongvaux, A. et al. Development and function of human innate immune cells in a humanized mouse model. Nat Biotechnol 32, 364–372 (2014).

14. Hoogstad-van Evert, J.S. et al. Umbilical cord blood CD34+ progenitor-derived NK cells efficiently kill ovarian cancer spheroids and intraperitoneal tumors in NOD/SCID/IL2Rgnull mice. OncoImmunology 6, e1320630 (2017).

15. Chang, D.-K. et al. Anti-CCR4 monoclonal antibody enhances antitumor immunity by modulating tumor-infiltrating Tregs in an ovarian cancer xenograft humanized mouse model. OncoImmunology 5, e1090075 (2016).

16. Gitto, S.B. et al. An autologous humanized patient-derived-xenograft platform to evaluate immunotherapy in ovarian cancer. Gynecologic Oncology 156, 222–232 (2020).

17. Odunsi, A. et al. Fidelity of human ovarian cancer patient-derived xenografts in a partially humanized mouse model for preclinical testing of immunotherapies. Journal for ImmunoTherapy of Cancer 8, e001237 (2020).

18. McIntosh, B.E. et al. Nonirradiated NOD,B6.SCID Il2rgamma-/- Kit(W41/W41) (NBSGW) mice support multilineage engraftment of human hematopoietic cells. Stem cell reports 4, 171–180 (2015).

19. Rahmig, S. et al. Improved Human Erythropoiesis and Platelet Formation in Humanized NSGW41 Mice. Stem cell reports 7, 591–601 (2016).

20. Brown, R.B., Madrid, N.J., Suzuki, H. & Ness, S.A. Optimized approach for Ion Proton RNA sequencing reveals details of RNA splicing and editing features of the transcriptome. PLOS ONE 12, e0176675 (2017).

21. Brayer, K.J., Frerich, C.A., Kang, H. & Ness, S.A. Recurrent Fusions in MYB and MYBL1 Define a Common, Transcription Factor-Driven Oncogenic Pathway in Salivary Gland Adenoid Cystic Carcinoma. Cancer Discov 6, 176–187 (2016).

22. Davis, M.P.A., van Dongen, S., Abreu-Goodger, C., Bartonicek, N. & Enright, A.J. Kraken: A set of tools for quality control and analysis of high-throughput sequence data. Methods 63, 41–49 (2013).

23. Wood, D.E. & Salzberg, S.L. Kraken: ultrafast metagenomic sequence classification using exact alignments. Genome Biology 15, R46 (2014).

24. Subramanian, A. et al. Gene set enrichment analysis: A knowledge-based approach for interpreting genome-wide expression profiles. Proceedings of the National Academy of Sciences 102, 15545–15550 (2005).

25. Giuntoli, R.L. et al. Ovarian Cancer-associated Ascites Demonstrates Altered Immune Environment: Implications for Antitumor Immunity. Anticancer Research 29, 2875–2884 (2009).

26. Wertel, I. et al. Macrophage-derived chemokine CCL22 and regulatory T cells in ovarian cancer patients. Tumour Biol 36, 4811–4817 (2015).

27. Luo, D., Wan, X., Liu, J. & Tong, T. Optimally estimating the sample mean from the sample size, median, mid-range, and/or mid-quartile range. Stat Methods Med Res 27, 1785–1805 (2018).

28. Dong, R. et al. Histologic and molecular analysis of patient derived xenografts of high-grade serous ovarian carcinoma. J Hematol Oncol 9 (2016).

29. Coppin, E. et al. Enhanced differentiation of functional human T cells in NSGW41 mice with tissue-specific expression of human interleukin-7. Leukemia 35, 3561–3567 (2021).

30. Milliken, D., Scotton, C., Raju, S., Balkwill, F. & Wilson, J. Analysis of chemokines and chemokine receptor expression in ovarian cancer ascites. Clinical cancer research : an official journal of the American Association for Cancer Research 8, 1108–1114 (2002).

31. Negus, R.P., Stamp, G.W., Hadley, J. & Balkwill, F.R. Quantitative assessment of the leukocyte infiltrate in ovarian cancer and its relationship to the expression of C-C chemokines. Am J Pathol 150, 1723–1734 (1997).

32. Bronger, H. et al. CXCL9 and CXCL10 predict survival and are regulated by cyclooxygenase inhibition in advanced serous ovarian cancer. Br J Cancer 115, 553–563 (2016).

33. Lieber, S. et al. Prognosis of ovarian cancer is associated with effector memory CD8+ T cell accumulation in ascites, CXCL9 levels and activation-triggered signal transduction in T cells. OncoImmunology 7, e1424672 (2018).

34. Sato, E. et al. Intraepithelial CD8+ tumor-infiltrating lymphocytes and a high CD8+/regulatory T cell ratio are associated with favorable prognosis in ovarian cancer. Proc Natl Acad Sci U S A 102, 18538–18543 (2005).

35. Kzhyshkowska, J. et al. Role of tumor associated macrophages in tumor angiogenesis and lymphangiogenesis. Frontiers in Physiology 5 (2014).

36. Lee, Y.C. et al. Malignant Bowel Obstruction in Advanced Gynecologic Cancers: An Updated Review from a Multidisciplinary Perspective. Obstet Gynecol Int 2018, 1867238 (2018).

37. Colvin, E.K. Tumor-associated macrophages contribute to tumor progression in ovarian cancer. Front Oncol 4, 137 (2014).

38. Kinjyo, I., Bragin, D., Grattan, R., Winter, S.S. & Wilson, B.S. Leukemia-derived exosomes and cytokines pave the way for entry into the brain. Journal of Leukocyte Biology 105, 741–753 (2019).

39. Rosato, R.R. et al. Evaluation of anti-PD-1-based therapy against triple-negative breast cancer patient-derived xenograft tumors engrafted in humanized mouse models. Breast Cancer Research 20, 108 (2018).

40. Ibrahim, A.M. et al. Diverse Macrophage Populations Contribute to the Inflammatory Microenvironment in Premalignant Lesions During Localized Invasion. Frontiers in Oncology 10 (2020).

41. Yoshimura, T. et al. Induction of Monocyte Chemoattractant Proteins in Macrophages via the Production of Granulocyte/Macrophage Colony-Stimulating Factor by Breast Cancer Cells. Frontiers in Immunology 7 (2016).

42. Kampan, N.C. et al. Interleukin 6 Present in Inflammatory Ascites from Advanced Epithelial Ovarian Cancer Patients Promotes Tumor Necrosis Factor Receptor 2-Expressing Regulatory T Cells. Frontiers in Immunology 8 (2017).

43. Sato, T. et al. Interleukin 10 in the tumor microenvironment: a target for anticancer immunotherapy. Immunologic Research 51, 170–182 (2011).

44. Matte, I., Lane, D., Laplante, C., Rancourt, C. & Piché, A. Profiling of cytokines in human epithelial ovarian cancer ascites. Am J Cancer Res 2, 566–580 (2012).

45. Mustea, A. et al. Expression of IL-10 in patients with ovarian carcinoma. Anticancer Res 26, 1715–1718 (2006).

46. Furukawa, S. et al. MCP-1 promotes invasion and adhesion of human ovarian cancer cells. Anticancer Res 33, 4785–4790 (2013).

47. Jeong, M., Wang, Y.Y., Choi, J.Y., Lim, M.C. & Choi, J.H. CC Chemokine Ligand 7 Derived from Cancer-Stimulated Macrophages Promotes Ovarian Cancer Cell Invasion. Cancers (Basel) 13 (2021).

48. Wang, Y. et al. Interleukin-6 signaling regulates anchorage-independent growth, proliferation, adhesion and invasion in human ovarian cancer cells. Cytokine 59, 228–236 (2012).

49. Browning, L., Patel, M.R., Horvath, E.B., Tawara, K. & Jorcyk, C.L. IL-6 and ovarian cancer: inflammatory cytokines in promotion of metastasis. Cancer Manag Res 10, 6685–6693 (2018).

50. Rios-Doria, J., Stevens, C., Maddage, C., Lasky, K. & Koblish, H.K. Characterization of human cancer xenografts in humanized mice. Journal for immunotherapy of cancer 8, e000416 (2020).

