## Supplementary Table S2 for "Humanized NBSGW PDX Models of Disseminated Ovarian Cancer Recapitulate Key Aspects of the Tumor Immune Environment within the Peritoneal Cavity"

**Supplementary Table S2. Patient characteristics for the three PDX models**

| Patient ID# | Diagnosis | Age at diagnosis | Platinum sensitivity | BRCA mutation status | Survival time after surgery |
|---|---|---|---|---|---|
| **PDX3** | HGSOC Stage 3c | 49 | Sensitive | wildtype | survived 6.5 years, DOD |
| **PDX9** | HGSOC Stage 3B | 56 | Resistant | wildtype | survived 18 months, DOD |
| **PDX18** | HGSOC Stage 3C | 60 | Sensitive | wildtype | recurrence at 1 year, lost to follow up |

** DOD: dead of disease*
