## Supplementary Fig S1 for "Humanized NBSGW PDX Models of Disseminated Ovarian Cancer Recapitulate Key Aspects of the Tumor Immune Environment within the Peritoneal Cavity"

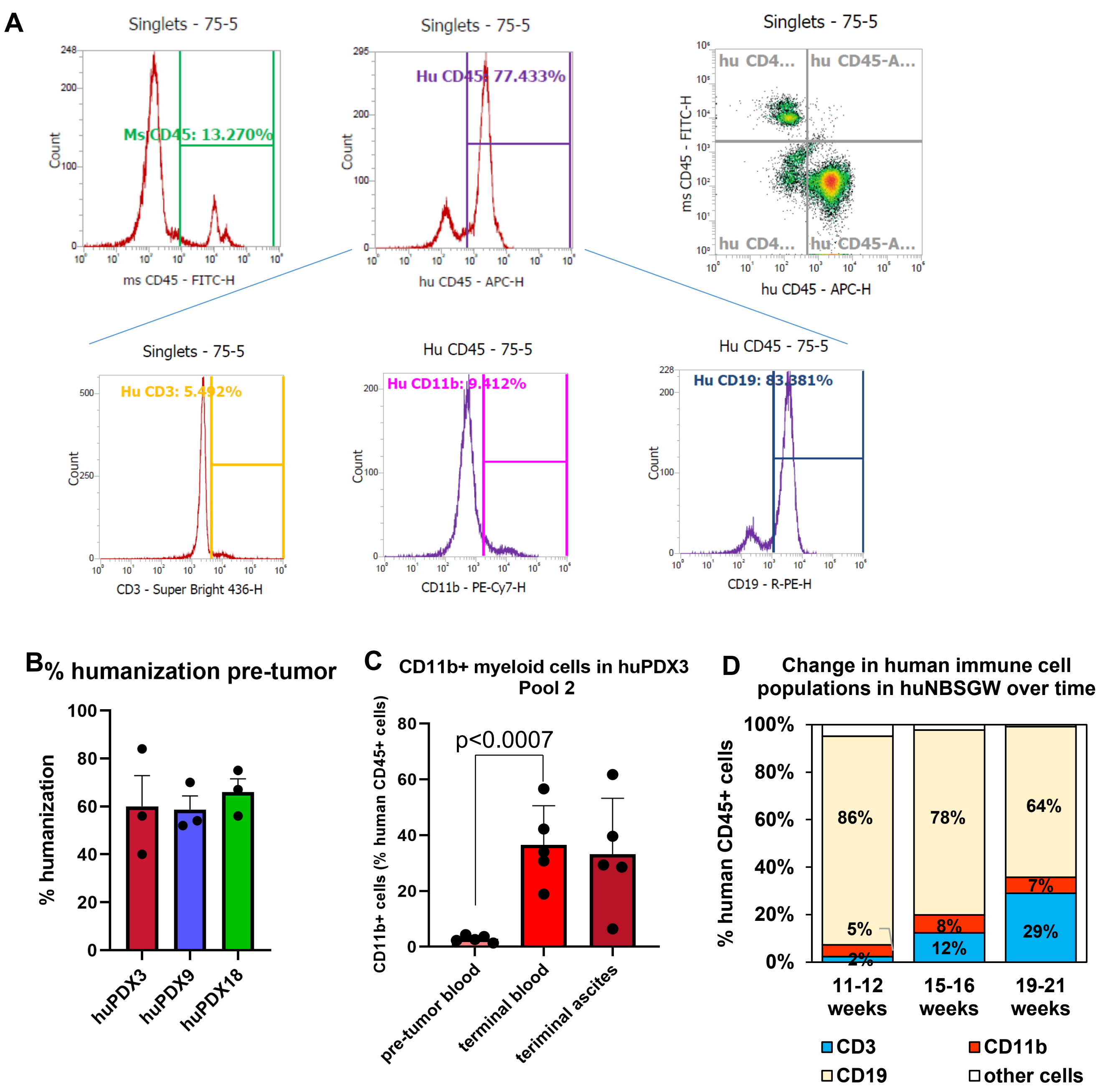


**Fig. S1 Analysis of humanization in mice engrafted with CD34+ hematopoietic stem cell.** Multi-color flow cytometry identified mouse and human CD45+ immune cells. Gating on human CD45+ blood cells, the human cells are then analyzed to determine the T cell (CD3+), B cell (CD19+) and myeloid cell (CD11b+) populations. **B.** Similar humanization in the peripheral blood from huNBSGW mice pre-PDX engraftment. The % humanization is the percent of human CD45+ cells in all CD45+ cells (mouse and human). There is no significant difference in humanization among the groups ($p<0.8$ by one-way ANOVA). **C**. Consistent increase in myeloid cells with tumor engraftment in huNBSGW PDX3. PDX3 (M-CSF high) demonstrates an increase in myeloid cells in peripheral blood at end point with a second independent donor pool of CD34+ cells. **D**. Change in the human immune profile of huNBSGW mice over time is marked by an increase in % human T-cells (CD3+) and a decrease in % human B-cells (CD19+). Myeloid cells (CD11b+) remain stable at 5-8%. The analysis is based on flow cytometry labeling of peripheral blood samples collected from huNBSGW at specified times post-engraftment.
