## Supplementary Fig S2 for "Humanized NBSGW PDX Models of Disseminated Ovarian Cancer Recapitulate Key Aspects of the Tumor Immune Environment within the Peritoneal Cavity"

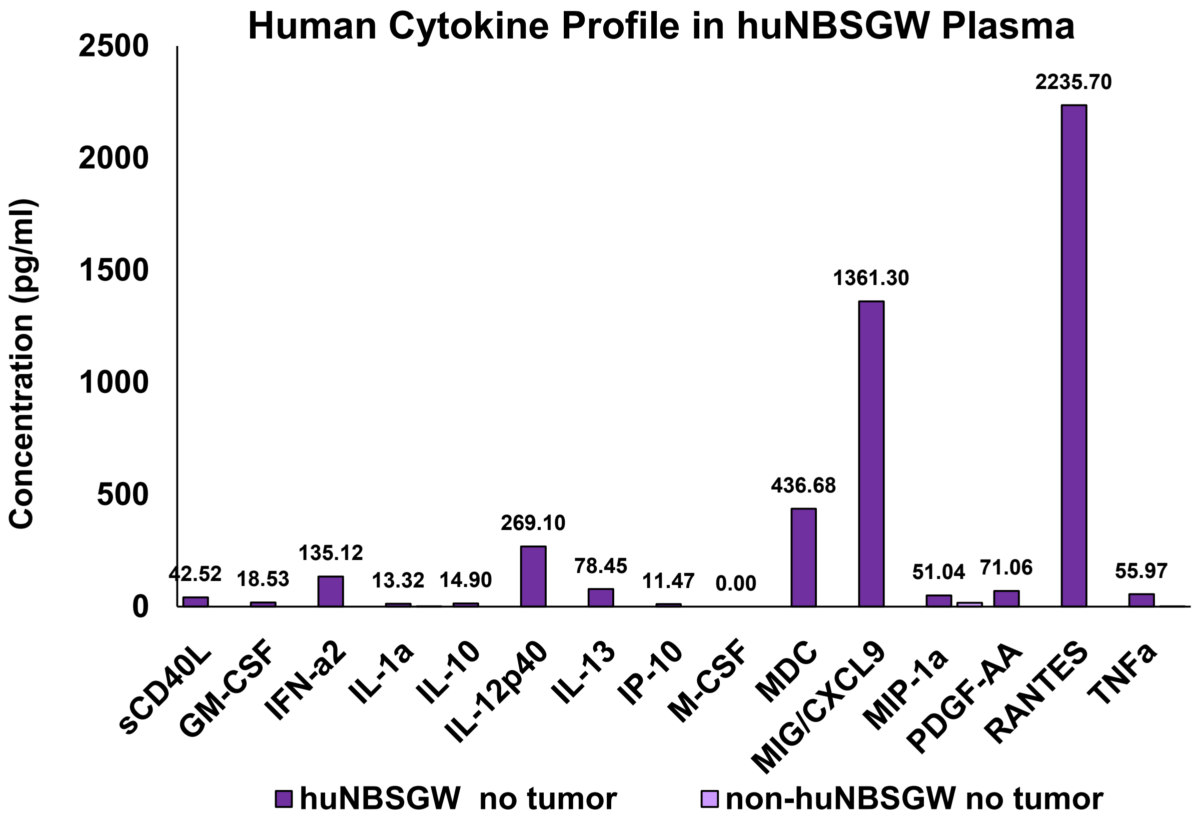


**Fig. S2. Levels of human factors in the plasma of a non-tumor bearing humanized NBSGW and a non-humanized non-tumor bearing NBSGW control.** Levels were measured on a 48-plex human cytokine/chemokine array.
