## Supplementary Table S3 for "Humanized NBSGW PDX Models of Disseminated Ovarian Cancer Recapitulate Key Aspects of the Tumor Immune Environment within the Peritoneal Cavity"

**Supplementary Table S3. Human cytokines with significantly higher levels in PDX ascites samples compared to plasma samples.** Factors are ordered based on ascites levels high to low. Levels are the log squares mean of all PDX ascites or plasma samples. Color scale shows relative levels with pink high and blue low.

| Cytokine | Ascites LSMeans | SEM | Plasma LSMeans | SEM | p-value |
|---|---|---|---|---|---|
| M-CSF | 8.2376 | 0.3515 | 2.9882 | 0.38212 | 0 |
| IP-10 | 7.975 | 0.60201 | 3.8294 | 0.65445 | 0.00011 |
| IL 8 | 7.6304 | 0.34784 | 5.0821 | 0.37815 | 0.00005 |
| MCP1 | 6.8919 | 0.54794 | 4.8412 | 0.59567 | 0.01856 |
| FGF_2 | 6.8143 | 0.30842 | 4.2355 | 0.33529 | 0.00001 |
| TNFα | 6.6726 | 0.32051 | 5.2466 | 0.34843 | 0.00621 |
| PDGF AA | 6.5517 | 0.41234 | 3.8245 | 0.44826 | 0.00017 |
| IL 27 | 6.3871 | 0.21342 | 4.2122 | 0.23201 | 0 |
| IL 1RA | 6.3749 | 0.55872 | 0.5132 | 0.6074 | 0 |
| MIG | 6.2657 | 0.21417 | 5.0478 | 0.23283 | 0.00082 |
| IL 6 | 6.2248 | 0.46911 | 4.0099 | 0.50998 | 0.00401 |
| IL 1a | 6.1278 | 0.4694 | 2.4501 | 0.51029 | 0.00002 |
| GM_CSF | 5.546 | 0.36579 | 3.1804 | 0.39766 | 0.00022 |
| Fractalkine | 5.5079 | 0.24343 | 2.884 | 0.26464 | 0 |
| IL 17E | 5.3828 | 0.37308 | 3.7895 | 0.40558 | 0.00823 |
| GROa | 5.3238 | 0.47464 | 2.9213 | 0.4366 | 0.00111 |
| IL 15 | 4.9463 | 0.23527 | 2.6377 | 0.25577 | 0 |
| TNFß | 4.9295 | 0.28047 | -1.1915 | 0.43575 | 0 |
| IL13 | 4.9114 | 0.4058 | -0.7923 | 0.44115 | 0 |
| IL 18 | 4.6658 | 0.34973 | 1.1234 | 0.3802 | 0 |
| IFNα2 | 4.266 | 0.25303 | 2.2373 | 0.27508 | 0.00002 |
| TGFα | 3.381 | 0.25799 | -0.9906 | 0.28047 | 0 |
| MIP 1ß | 3.3587 | 0.25445 | 2.2372 | 0.27661 | 0.00663 |
| MCP3 | 3.1507 | 0.35591 | 1.2868 | 0.38692 | 0.00173 |
| FLT3L | 2.8899 | 0.41323 | -1.1633 | 0.44923 | 0 |
| IL 1b | 2.8442 | 0.73826 | 0.0917 | 0.80258 | 0.01896 |
| sCD40L | 2.8064 | 0.20218 | 2.0481 | 0.2198 | 0.01834 |
| IL 4 | 2.6451 | 0.03536 | -1.2447 | 0.03844 | 0 |
| EGF | 1.7943 | 0.26219 | -0.8321 | 0.28503 | 0 |
| IL 17F | 1.7066 | 0.51958 | -0.7846 | 0.56484 | 0.00356 |
| Eotaxin | 1.6082 | 0.10024 | 0.6938 | 0.10897 | 0 |
| IL 7 | 1.2489 | 0.23602 | 0.2319 | 0.25658 | 0.00775 |
| IL 5 | 1.2036 | 0.29227 | 0.2869 | 0.31773 | 0.04469 |
| IL 2 | 0.7929 | 0.24787 | -0.6055 | 0.26946 | 0.00088 |
