## Supplementary Fig S3 for "Humanized NBSGW PDX Models of Disseminated Ovarian Cancer Recapitulate Key Aspects of the Tumor Immune Environment within the Peritoneal Cavity"

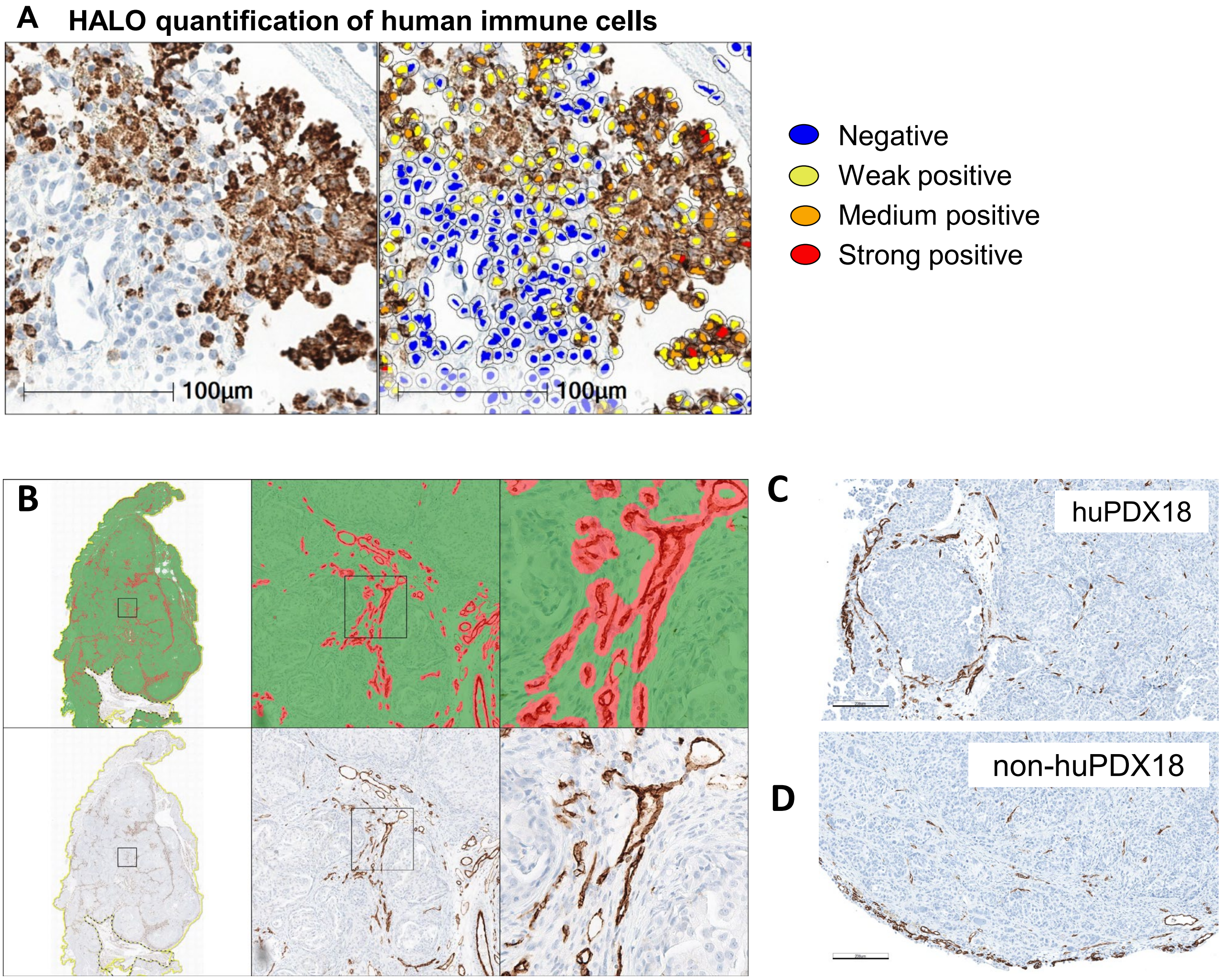


**Fig. S3. HALO Quantification of immune cells and vascular density. A**. An example of HALO quantification of human immune cells in a huPDX tumor. All cells within a tumor section are detected and then each cell is marked as negative or positive (weak, medium, strong) based on the analysis parameters. Scale bar=100 µm **B.** To quantify tumor vascularization in huPDX vs non-huPDX tumors, vessels were labeled with anti-CD31 antibody and vessel area was determined by analyzing scanned slides using the tissue classifier analysis software on the HALO platform. **C-D.** Comparison of vascularity in representative images from huPDX18 (**C**) and non-huPDX18 (**D**) labeled with anti-CD31 antibody. Scale bar=200 µm.
