## Supplementary Fig S4 for "Humanized NBSGW PDX Models of Disseminated Ovarian Cancer Recapitulate Key Aspects of the Tumor Immune Environment within the Peritoneal Cavity"

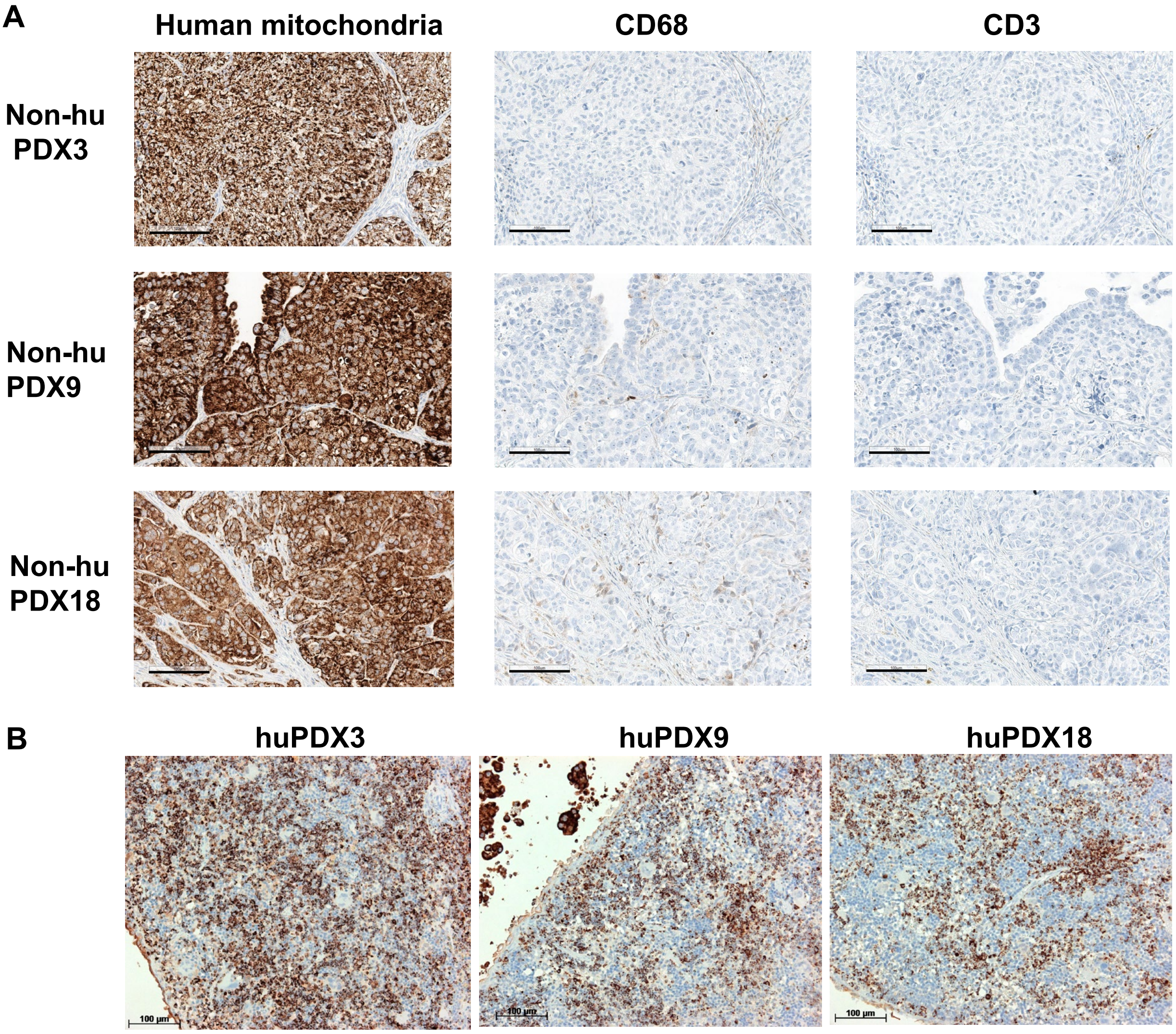


**Fig S4. Controls for human immune cell IHC A.** Immunohistochenmistry of non-huPDX control tumors. The human mitochondrial marker labels all PDX cancer cells. The human-specific markers for CD68 and CD3 show little to no background staining. **B.** Spleens from huPDX have populations of human immune cells labeled with the human mitochondria marker. Dark staining of cancer spheroids is also visible in the huPDX9 section (upper left). Scale bars=100 µm.
