## Supplementary Table S1 for "Humanized NBSGW PDX Models of Disseminated Ovarian Cancer Recapitulate Key Aspects of the Tumor Immune Environment within the Peritoneal Cavity"

### Supplementary Data

**Supplementary Table S1. Antibodies used for flow cytometry and IHC analysis.**

| **Antibody specificity** | **Cell type** | **Fluorescence** | **Antibody Clone** | **Supplier** | **Use** |
|---|---|---|---|---|---|
| Mouse CD45 | Mouse PBMCs | FITC | 30-F11 | BD Bioscience | Flow cytometry |
| Human CD45- | Human PBMCs | APC | HI30 | BD Bioscience | Flow cytometry |
| Human CD3 | Human T cells | BV421 | UCHT1 | BD Bioscience | Flow cytometry |
| CD11b | Human myeloid cells | PECy7 | ICRF44 | BD Bioscience | Flow cytometry |
| Human CD19 | Human B cells | PE | HIB19 | BD Bioscience | Flow cytometry |
| Human mitochondria | All human cells | | MAB1273 | Sigma | IHC |
| Human CD68 | Human TAMs | | PG-M1 | Agilent Dako | IHC |
| Human CD3 | Human T cells | | RBT-CD3 | Bio SB | IHC |
| Human CD4 | Human T cells | | SP35 | Sigma | IHC |
| Human CD8 | Human T cells | | SP57 | Ventana | IHC |
| Mouse CD31 | Mouse endothelial cells | | JC70A | Dako | IHC |
